# Increased prediction ability in Norway spruce trials using a marker x environment interaction and non-additive genomic selection model

**DOI:** 10.1101/478404

**Authors:** Zhi-Qiang Chen, John Baison, Jin Pan, Johan Westin, María Rosario García Gil, Harry X. Wu

## Abstract

A genomic selection (GS) study of growth and wood quality traits is reported based on control-pollinated Norway spruce families established in two Northern Swedish trials at two locations using exome capture as a genotyping platform. Non-additive effects including dominance and first-order epistatic interactions (including additive by additive, dominance by dominance, and additive by dominance) and marker-by-environment interaction (M×E) effects were dissected in genomic and phenotypic selection models. GS models partitioned additive and non-additive genetic variances more precisely compared with pedigree-based models. In addition, predictive ability (PA) in GS was substantially increased by including dominance and slightly increased by including M×E effects when these effects are significant. For velocity, response to GS (RGS) per year increased 91.3/43.7%, 86.9/82.9%, and 78.9/80.8% compared with response to phenotypic selection (RPS) per year when GS was based on 1) main marker effects (M), 2) M + M×E effects (A), and 3) A + dominance effects (AD) for site 1/site 2, respectively. This indicates that including M×E and dominance effects not only improves genetic parameter estimates but also may improve the genetic gain when they are significant. For tree height, Pilodyn, and modulus of elasticity (MOE), RGS per year improved up to 84.2%, 91.3%, and 92.6% compared with RPS per year, respectively.

## Introduction

Genomic selection (GS) is a breeding method that uses a dense set of genetic markers to accurately predict the genetic merit of individuals (Meuwissen et al. 2001) and it has been incorporated into animal breeding for many years (Van Eenennaam et al. 2014). Simulated studies have also shown that including dominance could increase the predictive ability (PA) (Nishio and Satoh, 2014) and result in a higher genetic gain in crossbred population when the dominance variance and heterosis are large and over-dominance is present (Zeng et al. 2013). In livestock, accounting for dominance in GS has improved genomic evaluations of dairy cows for fertility and milk production traits (Aliloo et al. 2016). In tree species, GS studies have been implemented in several breeding programs, but these studies mostly focused on additive effects in several commercially important conifer species, such as loblolly pine (*Pinus taeda* L.), Maritime pine (*Pinus pinaster* Ait.), Norway spruce (*Picea abies* (L.) Karst.), white spruce (*Picea glauca* (Moench) Voss) and hardwood Eucalypt species (Chen et al. 2018; Resende et al. 2012a; Resende et al. 2012b; Tan et al. 2017). The non-additive contributions have also been estimated in several studies (Bouvet et al. 2016; de Almeida Filho et al. 2016; Gamal El-Dien et al. 2016; Muñoz et al. 2014; Tan et al. 2018).

Based on several recent studies, dominance and epistasis may be confounded with the additive effects in both pedigree-based relationship matrix models (Gamal El-Dien et al. 2018) and genomic-based relationship matrix models (Tan et al. 2018). In the conventional pedigree-based genetic analysis, estimates of different genetic components such as additive, dominance and epistatic variances need full-sib family structure or full-sib family structure plus clonally replicated test (Mullin and Park, 1992). For most tree species, only a few reliable estimates for the non-additive variation were reported based on pedigree-based relationship (Baltunis et al. 2007; Isik et al. 2005; Isik et al. 2003; Weng et al. 2008; Wu et al. 2008), especially for wood quality traits (Wu, 2018).

The significant G×E is commonly observed among the different deployment zones for growth traits in Norway spruce (Chen et al. 2014; Chen et al. 2017; Kroon et al. 2011). Literature also supports the importance of predicting non-additive effects including dominance and epistasis in tree breeding program (Wu et al. 2016) and in clonal forestry program (Wu 2018). In a previous study (Chen et al. 2018), we used two full-sib family trials to study GS efficiency based on additive effects and different sampling strategies. Here, we extend our study to examine non-additive genetic effect using the genomic matrix and explore maker-by-environment interaction (M×E) effects on GS. The aims of the study are to 1) estimate and compare the non-additive genetic variances estimated from the average numerator relationship A-matrix (i.e. the expected theoretical relationships) and the realized genomic relationship G-matrix (i.e. the observed relationships); 2) evaluate the PA from different M×E models; 3) assess the PA of the model including non-additive effects; 4) evaluate the change of the breeding values ranking when models include the non-additive and M×E effects; and 5) assess genetic gain per year when M×E and dominance effects were included in the GS and phenotypic selection (PS) models.

## Materials and methods

### Sampling of plant material and genotyping

A total of 1,370 individuals was selected from two 28-year-old control-pollinated (full-sib) progeny trials. The progeny trials consist of the same 128 families generated through a partial diallel mating design involving 55 parents originating from Northern Sweden. Progenies were raised in the nursery at Sävar, and the trials were established in 1988 by Skogforsk in Vindeln (64.30°N, 19.67°E, altitude: 325 m) and in Hädanberg (63.58°N, 18.19°E, altitude: 240 m).

A completely randomized design without designed pre-block was used in the Vindeln trial (site 1), which was divided into 44 post-blocks. Each rectangular block has 60 trees (6 ×10) with expected 60 families at a spacing of 1.5 m × 2 m. The same design was also used in the Hädanberg trial (site 2) with 44 post-blocks. But for the purpose of demonstration, there was an extra block with 47 plots, each plot with 16 trees (4×4) planted in site 2. Based on the spatial analysis, in the final model, the 47 plots were combined into two big post-blocks.

### Phenotyping

Tree height was measured in 2003 at the age of 17 years. Solid-wood quality traits including Pilodyn penetration (Pilodyn) and acoustic velocity (velocity) were measured in October 2016. Surrogate wood density trait was measured using Pilodyn 6J Forest (PROCEQ, Zurich, Switzerland) with a 2.0 mm diameter pin, without removing the bark. Velocity is highly related to microfibril angle (MFA) in Norway spruce (Chen et al. 2015) and was determined using Hitman ST300 (Fiber-gen, Christchurch, New Zealand). By combining the Pilodyn penetration and acoustic velocity, indirect modulus of elasticity (MOE) was estimated using the equation developed in Chen et al. (2015).

### Genotyping

Buds and the first-year fresh needles from 1370 control-pollinated progeny trees and their 46 unrelated parents were sampled and genotyped using the Qiagen Plant DNA extraction protocol (Qiagen, Hilden, Germany) and DNA quantification was performed using the Qubit^®^ ds DNA Broad Range Assay Kit, Oregon, USA. The 46 parents were sampled in a grafted archive at Skogforsk, Sävar (63.89°N, 20.54°E) and in a grafted seed orchard at Hjssjö (63.93°N, 20.15°E). Probe design and evaluation are described in Vidalis et al. (2018). Sequence capture was performed using the 40 018 probes previously designed and evaluated for the materials (Vidalis et al. 2018) and samples were sequenced to an average depth of 15x at an Illumina HiSeq 2500 platform. The details of SNPs calling, filtering, quality control and imputation for this data could be found in Chen et al. (2018). Finally, 116,765 SNPs were kept for downstream analysis.

### Variance component and heritability models

The variance components and breeding values (BVs) for the genotypes of each trait in the two trials were estimated by using the best linear unbiased prediction (BLUP) method in three univariate models that included either additive (A), both additive and dominance (AD) or additive, dominance and epistasis genetic effects (ADE) as below. In practice, pedigree-based models had only two models because the epistatic effect is not possible to estimate in full-sib progeny trials without replicates for each genotype.

### Pedigree-based models

For the model with additive effect only (ABLUP-A):

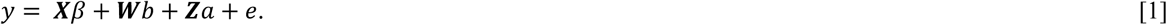

For the full model with both additive and dominance effects (ABLUP-AD):

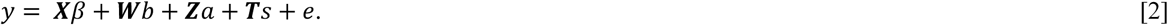

Where ***y*** is the vector of phenotypic observations of a single trait; *β* is the vector of fixed effects, including a grand mean and site effects, *b* is the vector of random post-block within site effects, *a* is the vector of random additive effects, *s* is the vector of random effects of specific combing ability (SCA), *e* is the random residual effects. ***X***, ***W***, ***Z,*** and ***T*** are the incidence matrices for *β*, *b*, a, and *s*, respectively. The random post-block effects (*b*) in equation [1] and [2] were assumed to follow 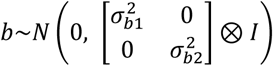, where *I* is the identity matrix, 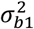 and 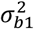 is the variance components of random post-block in site1 and site 2, respectively, and *⊗* is the Kronecker product operator. The random additive effects (*a*) were assumed to follow *a∼N(0, VCOV_a_ ⊗ A)*, where *A* is the pedigree-based additive genetic relationship matrix, *VCOV_a_* is the general case of additive variance and covariance structure. The random effects of SCA (*s*) were assumed to follow *s∼N(0, VCOV_s_ ⊗ I)*, where *VCOVs* is the general case of SCA variance and covariance structure. To study G×E in additive (*a*) and SCA (*s*) effects, six types of the different variance and covariance structures for additive effects including 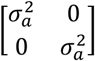 (IDEN), 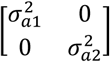 (DIAG), 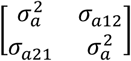 (CS), 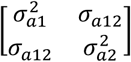 (CS+DIAG), 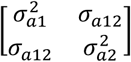 (US), and 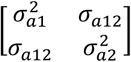 (FAMK) and for dominance effects including the same 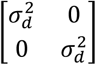 (IDEN), 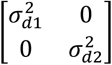 (DIAG), 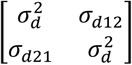 (CS), 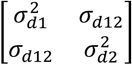 (CS+DIAG), 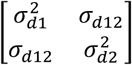 (US) and 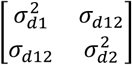 (FAMK) were used and represented in Table 1. 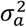 and 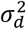 are the additive and dominance variances if homogenous variance structure were used 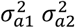, and *σ_a12_* are the additive variances for site 1, site 2 and additive covariance between site 1 and site 2, respectively. 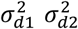, and *σ_d12_* are dominance variances for site 1, site 2 and additive covariance between site 1 and site 2. Our variance and covariances structures were similar to the published ones by Oakey et al. (2016). The US and FAMK structure are the same in any two sites MET model. The residual *e* was assumed to follow 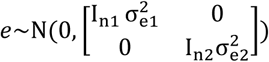, where 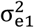 and 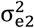 are the residual variances for site 1 and site 2, respectively, *I_n1_* and *I_n2_* are identity matrices, n1 and n2 are the number of individuals in each site. When the variance covariance structures (CS, CS+DIAG and FAMK) were used, the main marker/genetic effects (M) and M×E for additive, dominant, and epistatic variation could be predicted as described in Okay et al. (2016). For example, to estimate additive marker/genetic effects, *a* in equation [1] and [2] could be described as *a = m + me*, where *m* is the additive main marker/genetic effects, *me* is the additive main marker-by-environment effects.

**Table 1.** Six variance and covariance structures examined for the additive, dominance, and epistatic effects in two pedigree-based models and three genomic-based models.

| Structure | No. of parameter | Description |
| --- | --- | --- |
| IDEN | 1 | Identity |
| DIAG | n | Diagonal |
| CS | 2 | Compound symmetry |
| CS+DIAG | 1+n | Compound symmetry with heterogeneous variance |
| US | $n(n+1)/2$ | Unstructured |
| FAMK | $1+(k+1)n$ | Factor analytic with the main marker/genetic term and k factors |

n is the number of sites. k is the number of factors.

The pedigree-based additive (***A***) and dominance (***D***) relationship matrices were constructed based on the information from pedigree. The diagonal elements (*i*) of the *A* were calculated as *A_ii_ = 1 + f_i_ = 1 + A_gh_/2*, where *g* and *ℎ* are the *i*th individual’s parents, while the off-diagonal element is the relationship between individuals *i*th and *j*th calculated as *A_ij_ = A_ji_ = (A_jg_ + A_jh_)/2* (Mrode and Thompson, 2005). In the *D* matrix, the diagnonal elements were all one (*D_ii_ = 1*), while the off-diagonal elements between the individual *i*th and *j*th can be calculated as *D_ij_ = (A_gk_A_hl_ + A_gl_A_hk_)^⁄^4*, where *g* and *h* are the parents of *i*th’s individual and *k* and *l* are the parents of *j*th’s individual. *A* relationship matrix was produced using ASReml 4.1 (Gilmour et al. 2015) or ASReml-R package (Butler et al. 2009). *D* relationship matrix was produced using kin function in synbreed package in R (Wimmer et al. 2012).

### Genomic-based models

For the model with additive effect only (GBLUP-A):

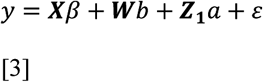

For the model with both additive and dominance effects (GBLUP-AD)

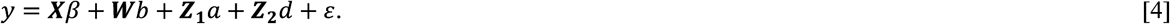

The full model (GBLUP-ADE) is described below:

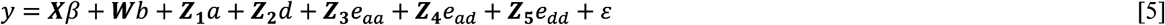

where *y* is the vector of phenotypic observations of a single trait; */3* is the vector of fixed effects, including the grand mean and site effects, *b* is the vector of random post-block within site effects following 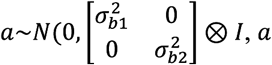 is the vector of random additive effects following *a∼N(0, VCOV_a_ ⊗ G_Ga_)*, *d* is the vector of random dominance effects following *d∼N(0, VCOV_d_ ⊗ G_d_)*, *e_aa_, e_ad_, and e_dd_* are the vectors of the random additive by additive epistatic effects, additive by dominance epistatic effects, and dominance by dominance epistatic effects, following 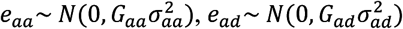, and 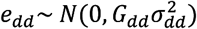, respectively. ***X***, ***W***, *Z_1_*, *Z_2_*, *Z_3_*, *Z_4_*, *and Z_5_* are the incidence matrices for *β*, *b*, a, *d*, *e_aa_*, *e_ad_* and *e_dd_*, respectively. *VCOV_Ga_* and *VCOV_Gd_* are the general cases of variance and covariance structures for additive and dominance effects, respectively. All variance covariance structures in Table 1 were used for genomic additive and dominance covariance structures in genomic-based models as in the pedigree-based models.

The genomic-based additive (*G_a_*) and dominance (*G_d_*) relationship matrices were constructed based on genome-wide exome capture data as described by VanRaden (2008) for *G_a_* and by Vitezica et al. (2013) for *G_d_*:

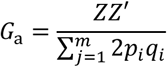

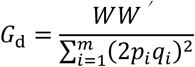

Where *m* is the total number of SNPs; the elements of *Z* are equal to *−2p_i_*, *q_i_ − p_i_*, and *2q_i_* for **aa**, **Aa**, and **AA** genotypes, respectively, with *p_i_* and *q_i_*being the allele frequency of **A** and **a** alleles at marker *i* in the population. For dominance matrix *G_d_*, **aa, Aa**, and **AA** genotypes in *W* were coded as 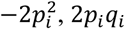, and 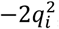, respectively. Based on the paper (Vitezica et al. 2013), the method guarantees the absence of confounding between *G_a_* and *G_d_* and could directly compare to the pedigree-based *A* and *D*.

The relationship matrices due to the first-order epistatic interactions were computed using the Hadamard product (cell by cell multiplication, denoted #) (Su et al. 2012). In the pedigree-based model, the additive by additive terms is based as *P_aa_ = A#A*, additive by dominance terms as *P_ad_ = A#D*, and dominance by dominance terms as *P_dd_ = D#D*. In genomic-based relationship matrices models: additive by additive terms *G_aa_ = G_a_#G_a_*, additive by dominance terms *G_ad_ = G_a_#G_d_*, and dominance by dominance terms *G_dd_ = G_d_#G_d_*.

Under the above models, the narrow-sense heritability can be estimated as 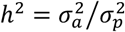, the dominance to total variance ratio as 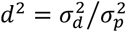, the epistatic to the total variance ratio as 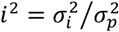 and the broad-sense heritability as 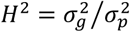, where 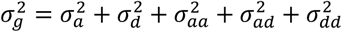 and 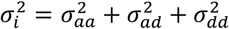. Broad-sense heritability for the ABLUP-AD model was estimated as 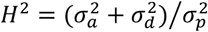 because epistatic effects were unable to be estimated.

### Model comparison

To compare the relative quality of the goodness-of-fit for the different models, Akaike Information Criterion (AIC) and the fitted line plot (graph of predicted 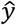 vs. adjusted y values) were used for the linear mixed-effects models (LMM) for all traits, while the standard error of the predictions (SEPs) of the traits BVs was used to assess the precision of the BVs.

### Cross-validation

A 10-fold cross-validation scenario with 10 replications was used to assess accuracy and prediction ability (PA).

### Expected performance of genomic selection

The expected performance of GS compared to the standard PS was evaluated only for GBLUP-AD model by calculating the response to genomic selection (RGS) as a percentage of the population average as follows:

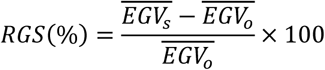

where 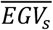 is the average of expected genetic values estimated from ABLUP-AD model for the selected population proportion based on 1) M, 2) M + M×E effects (A), or 3) A + dominance effects (AD) for site 1/site 2 from GBLUP-AD model, respectively, and 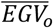 is the population average (Resende et al. 2017). The main advantage of GS is to shorten the breeding cycle. Thus, here we used RGS (%)/year and the response to phenotypic selection (RPS) (%)/year to compare the expected performances of GS and PS. In the Swedish Norway spruce breeding program, the traditional breeding cycle is at least 25 years long. If GS could be used as at a very early selection stage, the breeding cycle could be reduced to ca. 12.5 years based on the projection (Chen et al. 2018).

## Results

### Genetic variance components and heritability estimates

The six variance and covariance structures examined for the additive, dominance, and epistatic effects are presented in Table 1. The log-likelihood, Akaike Information Criterion (AIC) and Bayesian Information Criterion (BIC) for the five models (ABLUP-A, ABLUP-AD, GBLUP-A, GBLUP-AD, and GBLUP-ADE) under various variance structures are shown in Table S1 (supplementary data). The models with the best fitted variance-covariance structures under ABLUP and GBLUP for additive variance only, additive plus dominance variance or additive plus dominance and epistasis (e.g. ABLUP-A, ABLUP-AD, GBLUP-A, GBLUP-AD, and GBLUP-ADE) were listed in Table 2 and used to estimate the variance components (Table 3, except for Pilodyn with CS for additive effects and IDEN for dominance effects from GBLUP-A, GBLUP-AD, and GBLUP-ADE models). These models were included because we wanted to use the same variance-covariance structure to compare with the results from ABLUP-A and ABLUP-AD models for Pilodyn (Table 2).

**Table 2.** Summary of the five models (two ABLUP and three GBLUP models) with various variance and covariance structures fitted to the full data set for tree height, Pilodyn, velocity, and MOE

| Trait | Model | Variance structure |  |  |  | Log-likelihood | AIC | No. |
| --- | --- | --- | --- | --- | --- | --- | --- | --- |
|  |  | Additive | Dominance | Epistasis |  |  |  |  |
| Height | ABLUP-A | CS+DIAG |  |  |  | -6873.47 | 13760.95 | 7 |
|  | **ABLUP-AD** | **CS+DIAG** | **DIAG** |  |  | **-6868.92** | **13755.85** | **9** |
|  | GBLUP-A | CS+DIAG |  |  |  | -6874.05 | 13762.10 | 7 |
|  | **GBLUP-AD** | **CS+DIAG** | **IDEN** |  |  | **-6870.21** | **13756.42** | **8** |
|  | GBLUP-ADE | CS+DIAG | IDEN | IDEN-G3* |  | -6870.21 | 13762.42 | 11 |
| Pilodyn | **ABLUP-A** | **CS** |  |  |  | **-1727.77** | **3467.55** | **6** |
|  | ABLUP-AD | CS | IDEN |  |  | -1727.77 | 3469.55 | 7 |
|  | **GBLUP-A** | **IDEN** |  |  |  | **-1737.44** | **3484.88** | **5** |
|  | GBLUP-AD | IDEN | DIAG |  |  | -1735.87 | 3485.74 | 7 |
|  | GBLUP-ADE | IDEN | IDEN | IDEN-G3* |  | -1736.77 | 3493.54 | 10 |
| Velocity | ABLUP-A | CS |  |  |  | 1192.66 | -2373.33 | 6 |
|  | **ABLUP-AD** | **CS** | **IDEN** |  |  | **1194.59** | **-2375.19** | **7** |
|  | GBLUP-A | CS |  |  |  | 1183.37 | -2354.73 | 6 |
|  | **GBLUP-AD** | **CS** | **IDEN** |  |  | **1184.63** | **-2355.26** | **7** |
|  | GBLUP-ADE | CS | IDEN | IDEN-G3* |  | 1184.66 | -2349.32 | 10 |
| MOE | **ABLUP-A** | **CS** |  |  |  | **-2347.46** | **4706.92** | **6** |
|  | ABLUP-AD | CS | IDEN |  |  | -2347.46 | 4708.92 | 7 |
|  | **GBLUP-A** | **CS** |  |  |  | **-2357.84** | **4727.67** | **6** |
|  | GBLUP-AD | CS | IDEN |  |  | -2357.84 | 4729.67 | 7 |
|  | GBLUP-ADE | CS | IDEN | IDEN-G3* |  | -2357.84 | 4735.67 | 10 |

Variance and covariance structures: IDEN, identity; DIAG, Diagonal; CS, compound symmetry; CS+DIAG, compound symmetry with heterogeneous variance. \* G3 represents GBLUP-ADE model including three first order epistatic effects (the random additive by additive epistatic effects, additive by dominance epistatic effects, and dominance by dominance epistatic effects). No. is the number of variance parameters. Bold means the best model in GBLUP or ABLUP.

**Table 3.** Estimates of variance components (VC), their standard errors (SE) and the variance proportion of each site for tree height, Pilodyn, velocity, and MOE from the five genetic models fitted (ABLUP-A, ABLUP-AD, GBLUP-A, GBLUP-AD, and GBLUP-ADE)

| Trait | VC | ABLUP-A |  | ABLUP-AD |  | GBLUP-A |  | GBLUP-AD |  | GBLUP-ADE |  |
| --- | --- | --- | --- | --- | --- | --- | --- | --- | --- | --- | --- |
|  |  | Value (SE) | % | Value (SE) | % | Value (SE) | % | Value (SE) | % | Value (SE) | % |
| Height | $\sigma_{b_1}^2$ | 815.5 (330.2) | 12.1 | 804.0 (326.8) | 11.9 | 772.4 (317.8) | 11.5 | 703.1 (296.7) | 10.4 | 703.1 (296.7) | 10.4 |
| | $\sigma_{b_2}^2$ | 1962.0 (655.8) | 15.6 | 1863.3 (627.4) | 14.9 | 1916.2 (643.0) | 15.3 | 1918.6 (643.8) | 15.3 | 1918.6 (643.8) | 15.3 |
| | $\sigma_{a_1}^2$ | 690.8 (368.4) | 10.2 | 606.7 (392.1) | 9.0 | 902.2 (395.4) | 13.4 | 778.0 (407.2) | 11.5 | 778.0 (407.2) | 11.5 |
| | $\sigma_{a_{12}}^2$ | 565.9 (374.6) | | 571.2 (368.5) | | 573.1 (406.4) | | 463.4 (409.6) | | 463.4 (409.6) | |
| | $\sigma_{a_2}^2$ | 2007.1 (717.5) | 16.0 | 1371.5 (741.0) | 11.0 | 2140.6 (736.4) | 17.1 | 1858.7 (724.3) | 14.8 | 1858.7 (724.3) | 14.8 |
| | $\sigma_{d_1}^2$ | | | 572.2 (925.7) | 8.5 | | | 1224.1 (566.7) | 18.1 | 1224.1 (566.7) | 18.1 |
| | $\sigma_{d_2}^2$ | | | 2881.9 (1443.9) | 23.1 | | | 1224.1 (566.7) | 9.8 | 1224.1 (566.7) | 9.8 |
| | $\sigma_{aa}^2$ | | | | | | | | | 0.00 (0.00) | 0.0 |
| | $\sigma_{ad}^2$ | | | | | | | | | 0.00 (0.00) | 0.0 |
| | $\sigma_{dd}^2$ | | | | | | | | | 0.00 (0.00) | 0.0 |
| | $\sigma_{e_1}^2$ | 5260.5 (421.9) | 77.7 | 4777.6 (862.6) | 70.7 | 5064.1 (416.8) | 75.1 | 4053.3 (572.4) | 60.0 | 4053.28 (572.5) | 60.0 |
| | $\sigma_{e_2}^2$ | 8604.5 (640.7) | 68.4 | 6353.1 (1208.3) | 50.9 | 8461.8 (679.1) | 67.6 | 7523.9 (770.0) | 60.1 | 7523.86 (770.1) | 60.1 |
| | $h_1^2$ | 0.12 (0.06) | | 0.10 (0.06) | | 0.15 (0.06) | | 0.13 (0.06) | | 0.13 (0.06) | |
| | $h_2^2$ | 0.19 (0.06) | | 0.14 (0.07) | | 0.20 (0.06) | | 0.18 (0.06) | | 0.18 (0.06) | |
| | $H_1^2$ | | | 0.20 (0.14) | | | | 0.33 (0.09) | | 0.33 (0.09) | |
| | $H_2^2$ | | | 0.40 (0.12) | | | | 0.29 (0.07) | | 0.30 (0.07) | |
| Pilodyn | $\sigma_{b_1}^2$ | 0.24 (0.14) | 4.2 | 0.24 (0.14) | 4.2 | 0.23 (0.13) | 4.2 | 0.23 (0.13) | 4.2 | 0.23 (0.13) | 4.2 |
| | $\sigma_{b_2}^2$ | 0.63 (0.24) | 11.1 | 0.63 (0.24) | 11.1 | 0.71 (0.26) | 13 | 0.71 (0.26) | 13 | 0.71 (0.26) | 13.0 |
| | $\sigma_{a_1}^2$ | 2.30 (0.57) | 40.1 | 2.30 (0.57) | 40.1 | 1.79 (0.34) | 32 | 1.79 (0.34) | 32 | 1.79 (0.34) | 32.0 |
| | $\sigma_{a_{12}}^2$ | 2.03 (0.19) | | 2.03 (0.19) | | 1.61 (0.35) | | 1.61 (0.35) | | 1.61 (0.35) | |
| | $\sigma_{a_2}^2$ | 2.30 (0.57) | 40.7 | 2.30 (0.57) | 40.7 | 1.79 (0.34) | 32.8 | 1.79 (0.34) | 32.8 | 1.79 (0.34) | 32.8 |
| | $\sigma_{d_1}^2$ | | | 0 (0) | 0 | | | 0 (0) | 0 | 0 (0) | 0 |
| | $\sigma_{d_2}^2$ | | | 0 (0) | 0 | | | 0 (0) | 0 | 0 (0) | 0 |
| | $\sigma_{aa}^2$ | | | | | | | | | 0 (0) | 0 |
| | $\sigma_{ad}^2$ | | | | | | | | | 0 (0) | 0 |
| | $\sigma_{dd}^2$ | | | | | | | | | 0 (0) | 0 |
| | $\sigma_{e_1}^2$ | 3.20 (0.41) | 55.7 | 3.20 (0.41) | 55.7 | 0.32 (3.57) | 63.9 | 3.57 (0.32) | 63.9 | 3.57 (0.32) | 63.9 |
| | $\sigma_{e_2}^2$ | 2.72 (0.36) | 48.2 | 2.72 (0.36) | 48.2 | 0.27 (2.96) | 54.2 | 2.96 (0.27) | 54.2 | 2.96 (0.27) | 54.2 |
| | $h_1^2$ | 0.37 (0.09) | | 0.37 (0.09) | | 0.06 (0.30) | | 0.30 (0.06) | | 0.30 (0.06) | |
| | $h_2^2$ | 0.40 (0.09) | | 0.40 (0.09) | | 0.06 (0.34) | | 0.34 (0.06) | | 0.34 (0.06) | |
| | $H_1^2$ | | | 0.37 (0.09) | | | | 0.30 (0.06) | | 0.30 (0.06) | |
| | $H_2^2$ | | | 0.40 (0.09) | | | | 0.34 (0.06) | | 0.34 (0.06) | |
|  |  | Value (SE) | % | Value (SE) | % | Value (SE) | % | Value (SE) | % | Value (SE) | % |
| Velocity | $\sigma_{b_1}^2$ | 0.0018 (0.0013) | 2.4 | 0.0019 (0.1355) | 2.4 | 0.0013 (0.0011) | 1.8 | 0.0014 (0.0011) | 2.0 | 0.0014 (0.0011) | 2.0 |
| | $\sigma_{b_2}^2$ | 0.0036 (0.0018) | 4.6 | 0.0034 (0.2356) | 4.1 | 0.0033 (0.0017) | 4.4 | 0.0034 (0.0017) | 4.5 | 0.0034 (0.0017) | 4.5 |
| | $\sigma_{a_1}^2$ | 0.0365 (0.0087) | 48.1 | 0.0343 (0.0087) | 42.0 | 0.0305 (0.0051) | 42.4 | 0.0290 (0.0052) | 40.3 | 0.0282 (0.0065) | 39.1 |
| | $\sigma_{a_{12}}^2$ | 0.0320 (0.0086) | | 0.0293 (0.0087) | | 0.0241 (0.0051) | | 0.0224 (0.0052) | | 0.0215 (0.0066) | |
| | $\sigma_{a_2}^2$ | 0.0365 (0.0087) | 46.8 | 0.0343 (0.0087) | 40.9 | 0.0305 (0.0051) | 40.3 | 0.0290 (0.0052) | 38.3 | 0.0282 (0.0065) | 37.2 |
| | $\sigma_{d_1}^2$ | | | 0.0081 (0.0051) | 9.9 | | | 0.0067 (0.0046) | 9.3 | 0.0051 (0.0071) | 7.1 |
| | $\sigma_{d_2}^2$ | | | 0.0081 (0.0051) | 9.6 | | | 0.0067 (0.0046) | 8.8 | 0.0051 (0.0071) | 6.8 |
| | $\sigma_{aa}^2$ | | | | | | | | | 0.0030 (0.0152) | 4.2/4.0 |
| | $\sigma_{ad}^2$ | | | | | | | | | 0 (0) | 0 |
| | $\sigma_{dd}^2$ | | | | | | | | | 0.0005 (0.0108) | 0.7/0.7 |
| | $\sigma_{e_1}^2$ | 0.0376 (0.0057) | 49.5 | 0.0373 (0.0057) | 45.7 | 0.0402 (0.0042) | 55.8 | 0.0349 (0.0052) | 48.4 | 0.0336 (0.0077) | 46.7 |
| | $\sigma_{e_2}^2$ | 0.0379 (0.0053) | 48.6 | 0.0381 (0.0053) | 45.4 | 0.0418 (0.0040) | 55.3 | 0.0367 (0.0050) | 48.4 | 0.0355 (0.0079) | 48.1 |
| | $h_1^2$ | 0.43 (0.10) | | 0.40 (0.10) | | 0.34 (0.06) | | 0.32 (0.07) | | 0.31 (0.09) | |
| | $h_2^2$ | 0.43 (0.09) | | 0.39 (0.10) | | 0.33 (0.06) | | 0.31 (0.07) | | 0.30 (0.09) | |
| | $H_1^2$ | | | 0.51 (0.10) | | | | 0.41 (0.08) | | 0.43 (0.11) | |
| | $H_2^2$ | | | 0.50 (0.10) | | | | 0.40 (0.08) | | 0.42 (0.11) | |
| MOE | $\sigma_{b_1}^2$ | 0.49 (0.31) | 3.0 | 0.49 (0.31) | 3.0 | 0.45 (0.30) | 2.8 | 0.45 (0.30) | 2.8 | 0.45 (0.30) | 2.8 |
| | $\sigma_{b_2}^2$ | 0.68 (0.31) | 5.1 | 0.68 (0.31) | 5.1 | 0.87 (0.37) | 6.8 | 0.87 (0.37) | 6.8 | 0.87 (0.37) | 6.8 |
| | $\sigma_{a_1}^2$ | 6.63 (1.61) | 40.4 | 6.63 (1.61) | 40.4 | 5.23 (0.94) | 32.9 | 5.23 (0.94) | 32.9 | 5.23 (0.94) | 32.9 |
| | $\sigma_{a_{12}}^2$ | 5.82 (1.60) | | 5.82 (1.60) | | 4.54 (0.95) | | 4.54 (0.95) | | 4.54 (0.95) | |
| | $\sigma_{a_2}^2$ | 6.63 (1.61) | 50 | 6.63 (1.61) | 50 | 5.23 (0.94) | 41.1 | 5.23 (0.94) | 41.1 | 5.23 (0.94) | 41.1 |
| | $\sigma_{d_1}^2$ | | | 0 (0) | 0 | | | 0 (0) | 0 | 0 (0) | 0 |
| | $\sigma_{d_2}^2$ | | | 0 (0) | 0 | | | 0 (0) | 0 | 0 (0) | 0 |
| | $\sigma_{aa}^2$ | | | | | | | | | 0 (0) | 0 |
| | $\sigma_{ad}^2$ | | | | | | | | | 0 (0) | 0 |
| | $\sigma_{dd}^2$ | | | | | | | | | 0 (0) | 0 |
| | $\sigma_{e_1}^2$ | 9.29 (0.41) | 56.6 | 9.29 (1.17) | 56.6 | 10.24 (0.92) | 64.3 | 10.24 (0.92) | 64.3 | 10.24 (0.92) | 64.3 |
| | $\sigma_{e_2}^2$ | 5.96 (0.36) | 44.9 | 5.96 (0.95) | 44.9 | 6.61 (0.68) | 52.0 | 6.61 (0.68) | 52.0 | 6.61 (0.68) | 52.0 |
| | $h_1^2$ | 0.37 (0.09) | | 0.37 (0.09) | | 0.29 (0.05) | | 0.29 (0.05) | | 0.29 (0.05) | |
| | $h_2^2$ | 0.46 (0.10) | | 0.46 (0.10) | | 0.38 (0.07) | | 0.38 (0.07) | | 0.38 (0.07) | |
| | $H_1^2$ | | | 0.37 (0.09) | | | | 0.29 (0.05) | | 0.29 (0.05) | |
| | $H_2^2$ | | | 0.46 (0.10) | | | | 0.38 (0.07) | | 0.38 (0.07) | |

Notes: $\sigma_{b_1}^2$ and $\sigma_{b_2}^2$ are the block variance for site 1 and site 2. $\sigma_{a_1}^2$ , $\sigma_{a_2}^2$ , and $\sigma_{a_{12}}$ are the additive variances for site 1, site 2 and additive covariance between site 1 and site 2, respectively. $\sigma_{d_1}^2$ , $\sigma_{d_2}^2$ , and $\sigma_{d_{12}}$ are the dominance variances for site 1, site 2 and dominance covariance between site 1 and site 2. $\sigma_{aa}^2$ , $\sigma_{ad}^2$ , and $\sigma_{dd}^2$ are the additive × additive epistatic variance, additive × dominance epistatic variance, and dominance × dominance epistatic variances, respectively. $\sigma_{e_1}^2$ and $\sigma_{e_2}^2$ are the residual variances for site 1 and site 2, respectively. $h_1^2$ and $h_2^2$ are the narrow-sense heritability for site 1 and site 2, respectively. $H_1^2$ and $H_2^2$ are the broad-sense heritability for site 1 and site 2, respectively.

M×E effects for the additive or non-additive effects were considered significantly if the AIC value in MET analysis (e.g. under CS, CS+DIAG, US, or FAMK variance structure) was smaller than the corresponding AIC value in single site (ST) analysis (e.g. under IDEN or DIAG variance structure only) for the same trait or if the Log-likelihood Ratio test (LRT) was significant. All models with CS for additive genetic effects were found performing best, except for the model with CS+DIAG for tree height additive genetic effects (Table 2). Based on this criterion, all four traits were shown the significant additive M×E effects, except for the Pilodyn trait under GBLUP models. However, additive-by-environment variance in site 1 from ABLUP-AD with CS+DIAG was not significant (Table 3, 606.7) based on AIC. For dominance effect, however, only tree height with IDEN and velocity with DIAG structure had significant effects, therefore, there was no significant M×E for dominance effect for all traits. For epistasis, there was no significant effect on any traits.

Block variance components 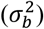 for each site were almost consistent across the five models (Table 3). For example, 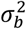 for tree height accounted for 10.4%−12.9% and 14.9%−15.6% for site 1 and site 2, respectively. For tree height, the main difference between the ABLUP-A and GBLUP-A models was the substantial increase of the additive variance 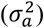 (Table 3), in contrast for wood quality traits. For example, tree height additive variance 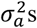 estimated from GBLUP-A were 130.6% and 106.7% of the ABLUP-A 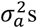 at site 1 and site 2, respectively. However, Pilodyn and velocity additive variance 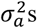 estimated from GBLUP-A averaged 77.8% and 83.6% of the ABLUP-A 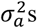 for both sites. The tree height 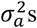 estimated from GBLUP-AD were also larger than those from ABLUP-AD for both sites. In contrast, wood quality traits 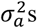 estimated from GBLUP-AD were also smaller than those from ABLUP-AD for both sites. For tree height and velocity, the main differences between the ABLUP-A and ABLUP-AD and between GBLUP-A and GBLUP-AD were the substantial decrease in 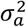 (Table 3). Pilodyn and MOE had the same 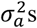 for the ABLUP-A and ABLUP-AD and also for GBLUP-A and GBLUP-AD because dominance variances 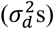 were zero for both traits. For example, tree height 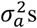 estimated from ABLUP-AD were 87.8% and 68.3% of the 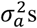 estimated from ABLUP-A at site 1 and site2, respectively.

In the ABLUP-AD model, tree height and velocity dominances showed significant effects based on AIC (Table 2 and Table 3). For example, tree height dominance effects accounted for 8.5% and 23.1% of the phenotypic variation for site 1 and site 2, respectively. In the GBLUP-AD model, tree height dominance effects accounted for 18.1% and 9.8% of the phenotypic variation for site 1 and site 2, respectively. However, the dominance effect of 572.2 in site 1 was not significant based on AIC. In the GBLUP-ADE models, all first-order epistatic effects were all zero for all the four traits, except for velocity with non-significant additive × additive effects (4.2%) and dominance × dominance effects (0.7%) (Table 3).

Tree height and velocity narrow-sense heritability estimates from ABLUP-A or GBLUP-A models were larger than those from ABLUP-AD or GBLUP-AD. For example, tree height narrow-sense heritability of 0.12 from ABLUP-A was larger than 0.10 from ABLUP-AD at site 1. Broad-sense heritability estimates were substantially larger than narrow-sense heritability estimates from both ABLUP-AD and GBLUP-AD at both sites for tree height and velocity. For example, tree height broad-sense heritability estimates were 253.8% and 166.7% of the narrow-sense heritability estimates from the GBLUP-AD model at site 1 and site 2, respectively. For tree height, Pilodyn, and MOE, GBLUP-ADE produced exactly the same results as GBLUP-AD (Table 3) because of lack of epistasis. In this study, only velocity showed non-significant and non-zero epistatic effects, moreover, broad-sense heritability estimates from GBLUP-ADE models were slightly higher than those from GBLUP-AD (0.43 vs. 0.41 for site 1, 0.42 vs. 0.40 for site 2).

### Comparison of models

We used two methods for model comparison, namely, Akaike Information Criterion (AIC) (Table S1 and Table 2) and the fitted line plots (represented by the graph of predicted values 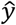 vs. observed values y) (Figure 1). The fitted line plot comparisons based on r^2^ reflected the goodness-of-fit. For tree height and velocity, r^2^ increased from GBLUP-A to GBLUP-AD (Tree height: 0.38 vs. 0.56 in site 1, 0.56 vs 0.79 in site 2; velocity: 0.80 vs. 0.88 in site 1, 0.78 vs. 0.87 in site 2) and from ABLUP-A to ABLUP-AD for both sites (Tree height: 0.44 vs. 0.74 in site 1, 0.58 vs 0.69 in site 2; velocity: 0.73 vs. 0.82 in site 1, 0.73 vs. 0.82 in site 2). For Pilodyn and MOE r^2^ was the same from GBLUP-A to GBLUP-AD and from ABLUP-A to ABLUP-AD, which was consistent with the zero estimates of dominance variances for both traits (Table 3). The difference of r^2^ for tree height between site 1 and site 2 was much larger than that of wood quality traits for all models.

**Figure 1.**
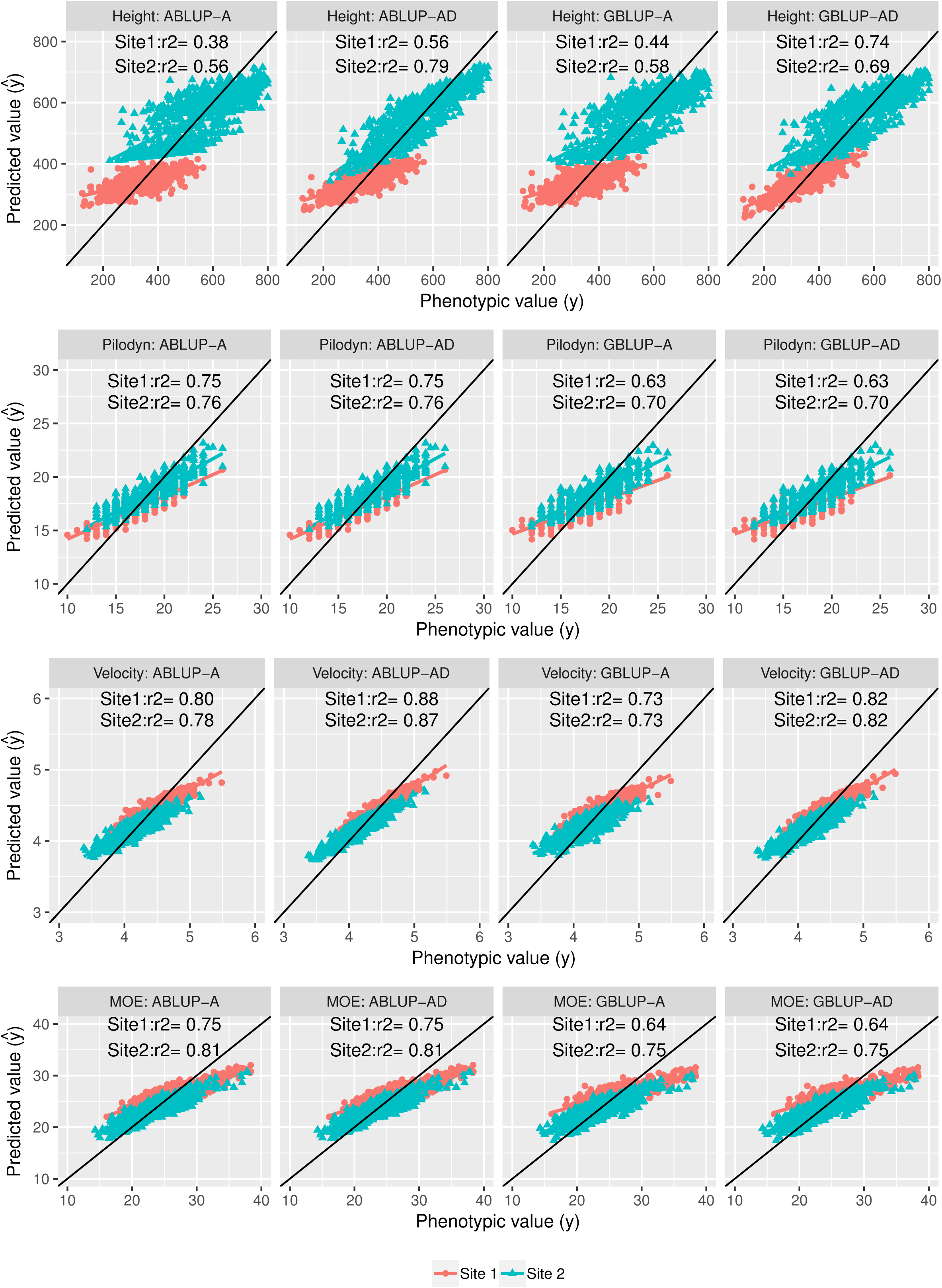
Model comparisons using the fitted line plots (represented by the graph of predicted values 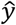 vs observed values y) for tree height, Pilodyn, velocity, and MOE.

Comparing BVs’ precision using the standard errors for the predictions (SEPs) between different modes (GBLUP-AD vs. GBLUP-A, GBLUP-AD vs. ABLUP-AD, GBLUP-AD vs. ABLUP-A, GBLUP-A vs. ABLUP-AD, GBLUP-A vs. ABLUP-A, and ABLUP-AD vs. ABLUP-A) is shown in Figure 2 for all traits. For tree height, the SEPs of 21-year-old Norway spruce breeding values between ABLUP-AD and ABLUP-A showed similar values. GBLUP-AD for tree height had much lower SEPs than that of GBLUP-A, but not for wood quality traits. GBLUP-AD for all traits had much lower SEPs values than that from ABLUP-AD for most SEPs values. ABLUP-AD for all traits had almost the same SEPs as ABLUP-A, even for tree height. For all traits, GBLUP-AD and GBLUP-A had more and lower SEPs than those from ABLUP-AD and ABLUP-A, except the GBLUP-A for tree height had more and larger SEPs than those from ABLUP-A and ABLUP-AD.

**Figure 2.**
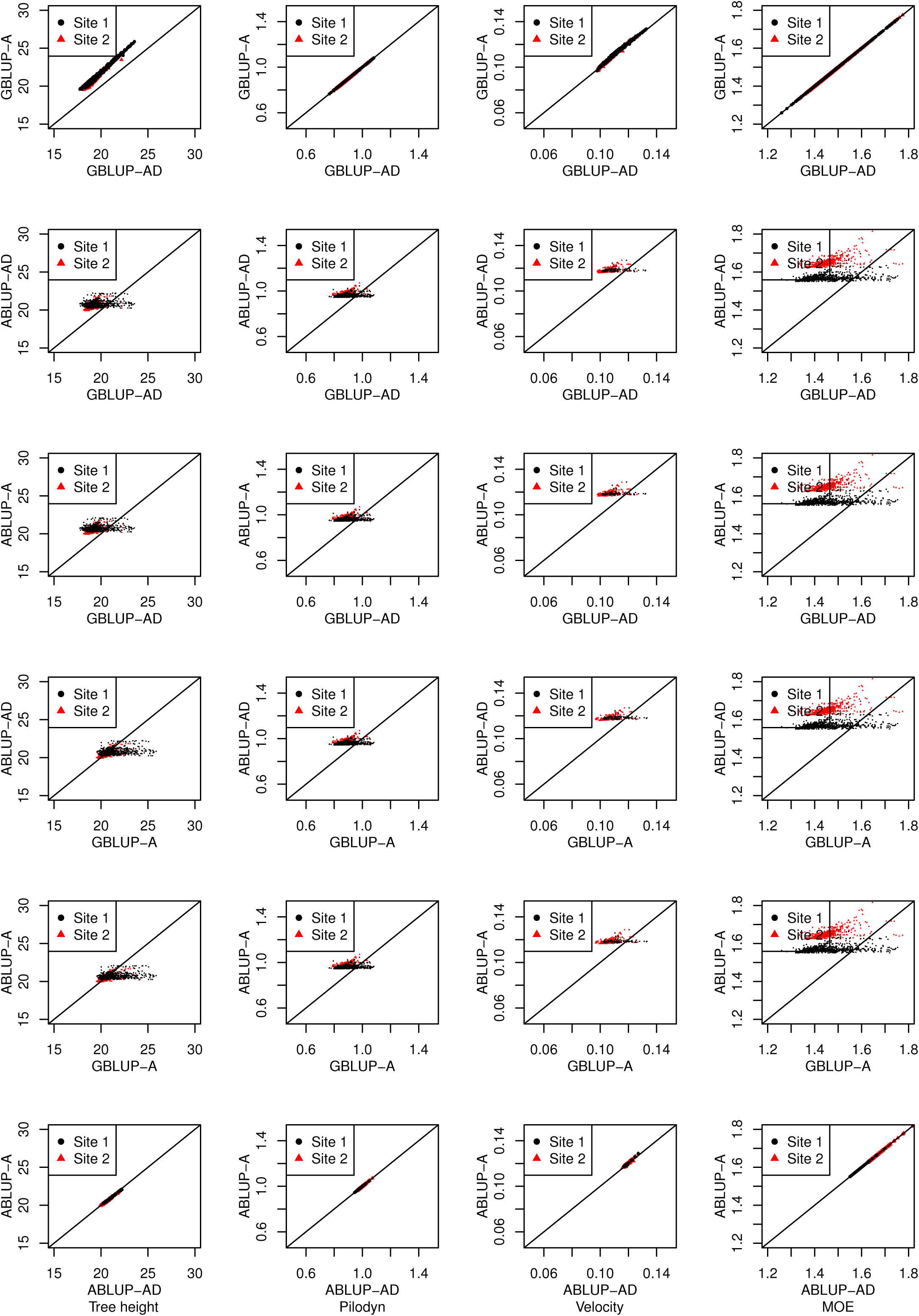
Standard errors for the predictions (SEP) for breeding values comparisons between GBLUP-AD and GBLUP-A, between GBLUP-AD and ABLUP-AD, between GBLUP-AD and ABLUP-A, between GBLUP-A and ABLUP-AD, between GBLUP-A and ABLUP-A, and between ABLUP-AD and ABLUP-A for tree height, Pilodyn, velocity, and MOE

### Cross-validation of the models

A random selection of 10% of the population was used as a validation set. Cross-validation was conducted using the same model for model building and validation and using different models for model building and validation (Table 4). Generally, using the same models have higher predictive accuracy than using different models (Table 4; diagonal values using the same models). Predictive accuracy was almost the same or similar among the pedigree-based models (between ABLUP-A and ABLUP-AD) and among the genomic-based models (between GBLUP-A and GBLUP-AD) (Table 4). For instance, predictive accuracies for Pilodyn between reference ABLUP-A and ABLUP-AD values with cross-validated ABLUP-A and ABLUP-AD values had all the same value (0.77). The predictive accuracy from ABLUP-AD was the highest (0.91).

**Table 4.**
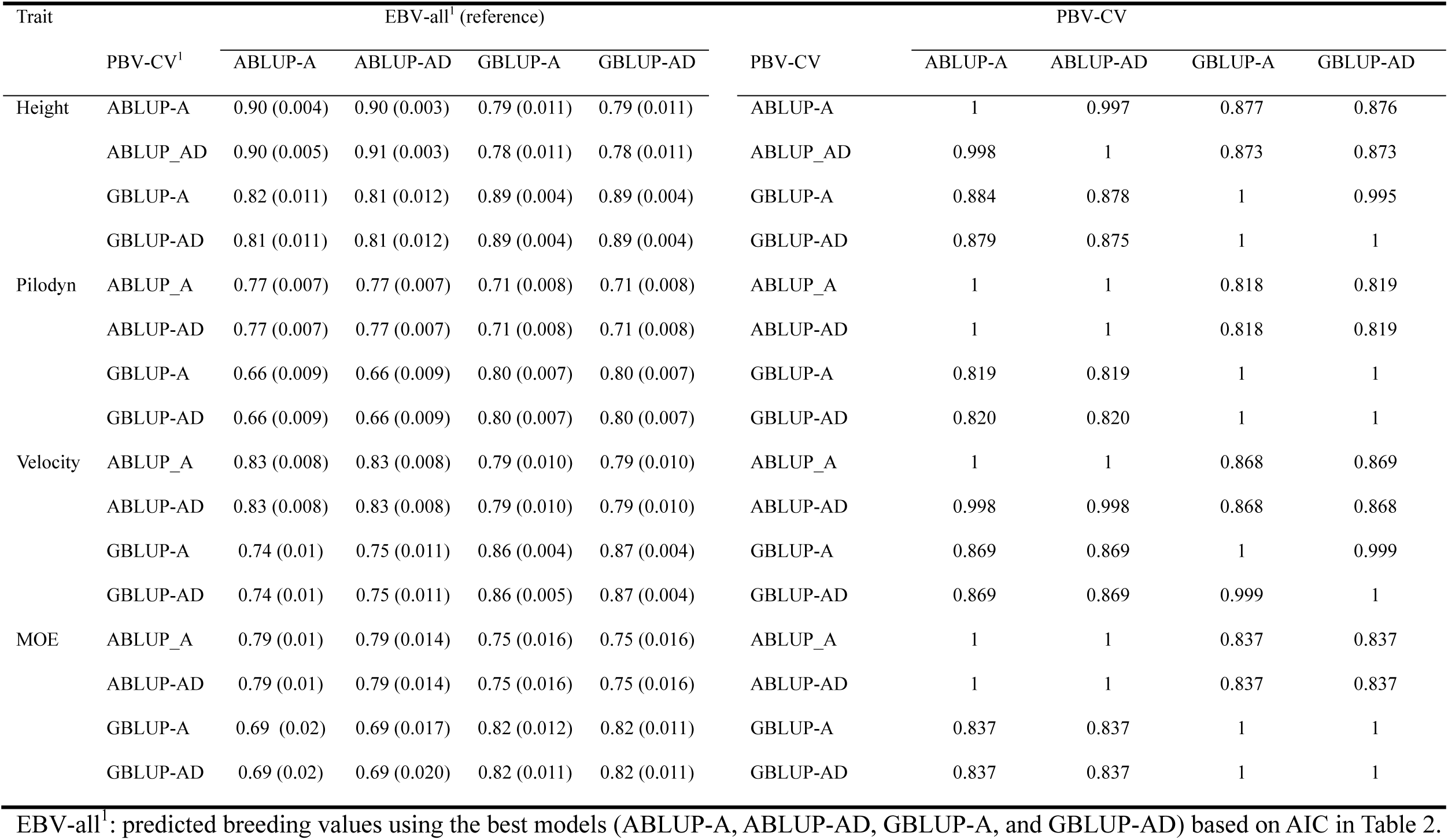
Pearson product-moment breeding value correlations between predicted breeding values using cross-validation (PBV-CV) and predicted breeding values (EBV-all)^1^ using the best models and Spearman breeding value ranking correlations between PBV-CV and PBV-CV within and among four models (ABLUP-A, ABLUP-AD, GBLUP-A, and GBLUP-AD) in cross-validation for tree height, Pilodyn, velocity, and MOE

For all four traits, Spearman breeding values ranking correlations between pedigree-based models (ABLUP-A and ABLUP-AD) and between genomic-based models (GBLUP-A and GBLUP-AD) in cross-validation were higher than between pedigree-based and genomic-based models (Table 4). For example, Spearman breeding values ranking correlation between pedigree-based and genomic-based models for tree height were 0.88. Spearman breeding values ranking correlations within pedigree-based models or genomic-based models were almost the same or unit. For example, Spearman breeding values ranking correlation between ABLUP-A and ABLUP-AD for tree height were 1.00.

The cross-validation focused on comparing the GBLUP and ABLUP models in prediction ability (PA) using MET models and single trial (ST) models for all traits and results were shown in Table 5. We only examined the models with either CS or CS+DIAG for additive effects and either CS or IDEN for dominance effects in MET analysis. For a single trial (ST) analysis, the models with DIAG for additive and IDEN or DIAG for dominance effects based on Table 2 were used. Using the same site data as training set and validation set showed higher PA. Tree height PA from the ST analysis in site 2 was higher than that in site 1 for additive effects (A) from GBLUP-AD models (comparisons: 1 and 3, 0.25 vs. 0.24, Table 5) and ABLUP-AD models (comparisons: 1 and 3, 0.26 vs. 0.21, Table 5). The models with additive and dominance effects (AD) showed similar results as the models with additive effect only (A) for tree height. If one site was used to building the model and predict the breeding values (A) and genotype values (AD) for the second site, then predicting for site 2 using the models from site 1 had a higher PA than the opposite for both GBLUP-AD (comparisons: 2 and 4, 0.09 vs. 0.07, Table 5) and ABLUP-AD (comparisons: 2 and 4, 0.13 vs. 0.09, Table 5). Ly et al. (2013) suggested that G×E, which can not be accounted for a single trial, reduced the ability to make predictions. Our results proved that the site 2 tree height might have a higher environmental component than those observed in site 1, making the prediction of the BVs (additive) or GVs (additive and dominance) less accurate. PA of Pilodyn did not change or only a slight change using site 1 model for site 2 and vice-versus. This happened because there is almost no G×E in Pilodyn measurement.

**Table 5.** Predictive abilities (PAs) based on main marker effects (M), M + marker-by-environment interaction effects (A) and A + dominance effects (AD) from GBLUP-AD and ABLUP-AD models for tree height, Pilodyn, velocity, and MOE in the single trial (ST) and multi-environment trial (MET) model analysis in cross-validation

| Trait | Comparison | Model | Structure | Training | Validation | GBLUP-AD |  |  | ABLUP-AD |  |  |
| --- | --- | --- | --- | --- | --- | --- | --- | --- | --- | --- | --- |
|  |  |  |  |  |  | M | A | AD | M | A | AD |
| Height | 1 | ST | $M_{DIAG}$ | Site 1 | Site1 | N/A | 0.24 (0.04) | 0.26 (0.03) | N/A | 0.21 (0.04) | 0.20 (0.04) |
| | 2 | ST | $M_{DIAG}$ | Site 1 | Site2 | N/A | 0.09 (0.03) | 0.16 (0.03) | N/A | 0.13 (0.03) | 0.12 (0.03) |
| | 3 | ST | $M_{DIAG}$ | Site 2 | Site2 | N/A | 0.25 (0.03) | 0.27 (0.03) | N/A | 0.26 (0.04) | 0.29 (0.04) |
| | 4 | ST | $M_{DIAG}$ | Site 2 | Site1 | N/A | 0.07 (0.04) | 0.12 (0.04) | N/A | 0.09 (0.03) | 0.08 (0.03) |
| | 5 | MET | $M_{CS+DIAG}$ | Site 1 | Site 1 | 0.22 (0.04) | 0.23 (0.04) | 0.26 (0.03) | 0.19 (0.03) | 0.19 (0.03) | 0.20 (0.03) |
| | 6 | MET | $M_{CS+DIAG}$ | site 2 | Site 2 | 0.22 (0.04) | 0.25 (0.03) | 0.27 (0.03) | 0.21 (0.03) | 0.24 (0.03) | 0.29 (0.04) |
| Pilodyn | 1 | ST | $M_{DIAG}$ | Site 1 | Site 1 | N/A | 0.26 (0.05) | 0.27 (0.05) | N/A | 0.30 (0.05) | 0.30 (0.05) |
| | 2 | ST | $M_{DIAG}$ | Site 1 | Site 2 | N/A | 0.23 (0.04) | 0.23 (0.04) | N/A | 0.24 (0.03) | 0.25 (0.03) |
| | 3 | ST | $M_{DIAG}$ | Site 2 | Site 2 | N/A | 0.23 (0.03) | 0.31 (0.03) | N/A | 0.34 (0.02) | 0.33 (0.02) |
| | 4 | ST | $M_{DIAG}$ | Site 2 | Site 1 | N/A | 0.23 (0.03) | 0.23 (0.03) | N/A | 0.23 (0.03) | 0.24 (0.03) |
| | 5 | MET | $M_{CS}$ | Site 1 | Site 1 | 0.30 (0.04) | 0.30 (0.04) | 0.30 (0.04) | 0.32 (0.03) | 0.33 (0.04) | 0.33 (0.04) |
| | 6 | MET | $M_{CS}$ | Site 2 | Site 2 | 0.32 (0.03) | 0.32 (0.03) | 0.32 (0.03) | 0.34 (0.02) | 0.35 (0.02) | 0.35 (0.02) |
| Velocity | 1 | ST | $M_{DIAG}$ | Site 1 | Site 1 | N/A | 0.44 (0.04) | 0.45 (0.04) | N/A | 0.40(0.04) | 0.42 (0.04) |
| | 2 | ST | $M_{DIAG}$ | Site 1 | Site 2 | N/A | 0.32 (0.03) | 0.33 (0.02) | N/A | 0.35 (0.03) | 0.36 (0.03) |
| | 3 | ST | $M_{DIAG}$ | Site 2 | Site 2 | N/A | 0.38 (0.02) | 0.39 (0.02) | N/A | 0.40 (0.04) | 0.41 (0.04) |
| | 4 | ST | $M_{DIAG}$ | Site 2 | Site 1 | N/A | 0.34 (0.06) | 0.35 (0.06) | N/A | 0.36 (0.04) | 0.38 (0.04) |
| | 5 | MET | $M_{CS}$ | Site 1 | Site 1 | 0.45 (0.05) | 0.46 (0.04) | 0.46 (0.04) | 0.42 (0.04) | 0.43 (0.04) | 0.43 (0.04) |
| | 6 | MET | $M_{CS}$ | Site 2 | Site 2 | 0.39 (0.03) | 0.39 (0.03) | 0.39 (0.03) | 0.42 (0.04) | 0.43 (0.04) | 0.43 (0.04) |
| MOE | 1 | ST | $M_{DIAG}$ | Site 1 | Site 1 | N/A | 0.33 (0.03) | 0.33 (0.03) | N/A | 0.34 (0.03) | 0.35 (0.03) |
| | 2 | ST | $M_{DIAG}$ | Site 1 | Site 2 | N/A | 0.28 (0.04) | 0.28 (0.04) | N/A | 0.31 (0.04) | 0.32 (0.04) |
| | 3 | ST | $M_{DIAG}$ | Site 2 | Site 2 | N/A | 0.33 (0.03) | 0.33 (0.04) | N/A | 0.36 (0.04) | 0.36 (0.04) |
| | 4 | ST | $M_{DIAG}$ | Site 2 | Site 1 | N/A | 0.30 (0.04) | 0.30 (0.04) | N/A | 0.32 (0.04) | 0.32 (0.04) |
| | 5 | MET | $M_{CS}$ | Site 1 | Site 1 | 0.37 (0.04) | 0.37 (0.04) | 0.37 (0.04) | 0.38 (0.03) | 0.39 (0.03) | 0.39 (0.03) |
| | 6 | MET | $M_{CS}$ | Site 2 | Site 2 | 0.35 (0.04) | 0.35 (0.04) | 0.35 (0.04) | 0.38 (0.04) | 0.38 (0.04) | 0.38 (0.04) |

Generally, PA was higher in MET analysis than that in ST analysis for all traits, except for tree height (Table 5). For Pilodyn, velocity, and MOE, PAs in MET analyses based on A and AD models were higher than those from single site (ST) analyses (comparisons 1 and 5, comparisons 2 and 6, Table 5). For example, PAs for Pilodyn based on A from GBLUP-AD showed an increase of 15.4% (comparison1 1 and 5, 0.26 vs. 0.30, Table 5) and 39.1% (comparisons 3 and 6, 0.23 vs. 0.32, Table 5) in site1 and site 2, respectively.

Finally, we studied the additive M×E effects on the GEBVs. There was a reduction in tree height PA if M×E was not included in calculating the GEBVs for site 2 (comparison 6: 0.25 vs. 0.22, Table 5), and for site 1 (comparison 5: 0.23 vs. 0.22, Table 5). Including tree height dominance in models in site 2, PA increased 8% from 0.25 to 0.27 and 20.8% from 0.24 to 0.29 for GBLUP-AD and ABLUP-AD models, respectively (Table 5). Including tree height dominance in models in site 1, PA increased 13.0% from 0.23 to 0.26 and 5.3% from 0.19 to 0.20 for GBLUP-AD and ABLUP-AD models, respectively. For Pilodyn, velocity, and MOE, PA including dominance in MET analysis were not increased, even for velocity with a significant dominance variance based on AIC.

PAs for all traits from GBLUP-ADE were not shown in Table 5 because their variance components were zero, except for velocity. PA for velocity from the GBLUP-ADE model was the same as the result from GBLUP-AD.

### Expected response to genomic selection (GS)

We compared GS at seedling stage (about age two years) with the phenotypic selection at age ca. 30 years (i.e. a minimum of 25 years), as in the traditional breeding program in Northern Sweden. A conservative response of genomic selection per year (RGS%/year) was calculated to compare with the response of phenotypic selection per year (RPS%/year) for variable proportions of individuals selected by GS. We compared RGS per year with RPS per year for all traits for variable proportions of individuals selected by GS (Figure 3). The results showed that RGS per year provided much larger values than RPS per year for three genomic selection scenarios, including selection based on 1) main marker effects (M), 2) M plus M×E effects (A), and 3) A plus dominance effects (AD) from GBLUP-AD model for both sites (Figure 3). However, RGS per year for different scenarios in both sites showed slight differences only for tree height, not for wood quality traits. RGS per year for tree height based on A and AD was substantially higher than that based on M in site 2 (Figure 3). However, in site 1, RGS per year for tree height based on A and AD was slightly better than that based on M and only showed at low selection proportion. In the traditional Swedish breeding program, 50 individuals were selected for each breeding population. In Figure S1, the top 50 individuals selected without considering relationships for selection based on M, A, and AD for all traits were scaled to the total expected genetic value (EGV) ranking of all individuals in site 1 and site 2. RGS per year based on M, A, and AD for GBLUP-AD and RPS per year based on an AD for ABLUP-AD are shown in Table S2. For tree height, RGS per year based on AD in site 2 was 0.54 (%)/year, which was substantially higher than 0.45 and 0.46 (%)/year based on A and M, respectively. RGS per year based on AD in site 1 was 0.43 (%)/year, which was slightly higher than 0.41 and 0.41 (%)/year based on A and M, respectively. For wood quality traits, RGS per year based on M, A, and AD were almost the same (Figure 3), but they slightly increased when such effects were significant (Table S2). If the top 50 velocity individuals based on GEGVs selected, RGS per year from GBLUP-AD were 91.3%, 86.9%, and 78.9% in site 1 and 88.2%, 82.9%, and 80.8% in site 2 higher than RPS (%)/year based on M, A, and AD effects, respectively. RGS per year from GBLUP-AD for tree height, Pilodyn, and MOE were up to 68.9%, 91.3%, and 92.6%, respectively.

**Figure 3.**
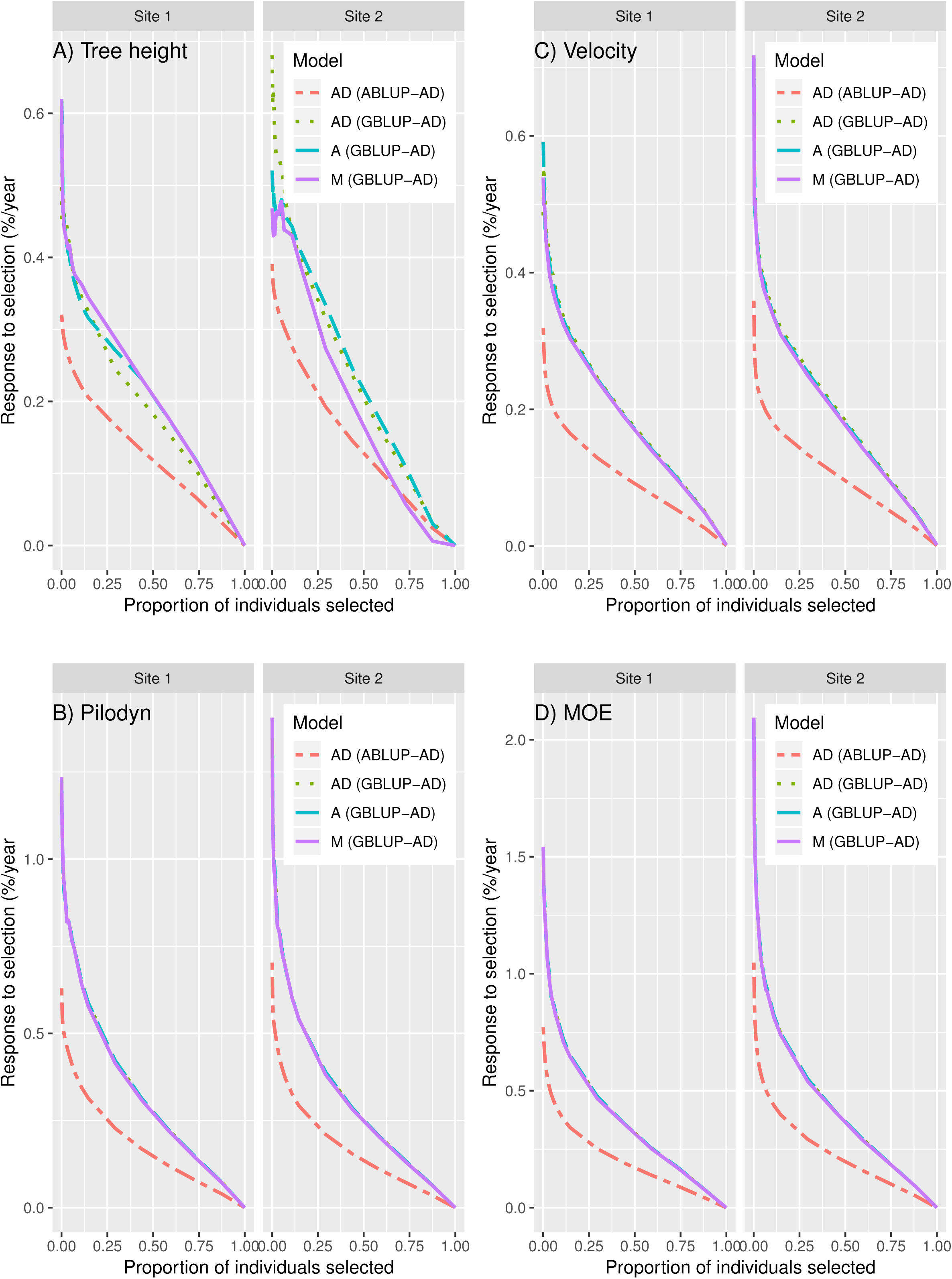
Response to genomic selection (RGS), including three different selection scenarios based on 1) only main marker effects (M), 2) main marker effects plus genotype-by-environment interaction effects (A), and 3) A + dominance (AD) from GBLUP-AD and ABLUP-AD models for A) tree height, B) Pilodyn, C) velocity, and D) MOE, expressed as a percentage gain of the average population mean per year, compared with response to phenotypic selection (RPS) also including dominance effect (ABLUP-AD) calculated for different proportions of individuals selected by GS.

## Discussion

### Genetic variance components and heritability estimates

In the traditional Norway spruce breeding program, estimates of broad-sense heritability (*H*^2^) have previously been conducted in clonal tests where the clones have been selected from commercial nurseries. For example, tree height *H*^2^ estimates vary from 0.12 to 0.44 for Norway spruce (Bentzer et al. 1989; Karlsson and Högberg 1998; Karlsson et al. 2000), but it is not possible to compare with narrow-sense heritability (*h*^2^), which requires family structure. Epistasis estimation, on the other hand, requires full-sib family structure plus the replication for genotypes in clonal trials. Existing high-throughput single nucleotide polymorphism (SNP) genotyping technology such as SNP arrays, re-sequencing or GBS allows genotyping for larger numbers of SNPs, and therefore uses to study dominance and epistasis in populations without pedigree delineation of full-sib family structure in animal (Aliloo et al. 2016; Sun et al. 2013) and plants (Gamal El-Dien et al. 2018).

In the previous study, it was observed that the average *h*^2^ of 0.29 (0.02–1.09) based on 170 field tests with seedling materials was higher than the average *H*^2^ of 0.18 (0.04–0.50) based on 123 field tests with clonal materials (Kroon et al. 2011), indicating a valid comparison of relative genetic control must use the datasets that come from the same trial with comparable pedigree (Wu, 2018). In this study, tree height *h*^2^ estimated from pedigree-based models (Table 3: ABLUP-A and ABLUP-AD) varies from 0.10 to 0.19. The tree height *H*^2^ estimated from pedigree-based models varies from 0.20 to 0.40 in the two sites. The ratio of tree height *h*^2^/*H*^2^ varies from 0.35–0.50 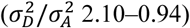 and from 0.39–0.60 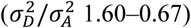 in ABLUP-AD and GBLUP-AD models, respectively, which are lower than 0.60–0.84 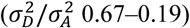 from three Norway spruce progeny trials in the previous study (Kroon et al. 2011). The usual range of the ratio *h*^2^/*H*^2^ has been reported to range from 0.18 to 0.84 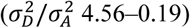 for tree traits (White et al. 2007; McKeand et al. 2008; Wu et al. 2008; Bian et al. 2014). It is also considered that significant dominance could be utilized in advanced Norway spruce breeding and deployment programs.

Our results show that the inclusion of dominance effects reduces the estimates of *h*^2^ from GBLUP-AD and ABLUP-AD when dominance is not zero. For example, tree height *h*^2^ estimate decreases 13%–26%, which is similar to hybrid *Eucalyptus* with a substantial decrease (50%–70%) reported by Tan et al. (2018). The situation is expected from a theoretical standpoint as a substantial proportion of the non-additive variance can be inseparable from additive variance (Falconer and Mackay, 1996), that has been encountered in the several empirical results (Gamal El-Dien et al. 2016; Gamal El-Dien et al. 2018; Tan et al. 2018).

### Comparison and cross-validation of models

AIC values for the GBLUP models were not significantly higher than those based on pedigree relationship matrices, which is consistent with the results of Gamal El-Dien et al. (2016) in white spruce, but is in contrast with the results from hybrid *Eucalyptus* (Tan et al. 2018). In the *Eucalyptus* study, the significant improvement from GBLUP models based on AIC may result from a considerable number of uncorrected wrong pedigrees. In our study, the SEPs of breeding values in GBLUP-A models for tree height was higher than that in pedigree-based ABLUP-A model (Figure 2), which is inconsistent with the results of Gamal El-Dien et al. (2018). This seems reasonable in our study because the additive variance increases from ABLUP models to GBLUP models (Table 3). For wood quality traits, the SEPs of breeding values in GBLUP-A models are smaller than those in the pedigree-based ABLUP-A model. For all traits, the most SEPs of breeding values in GBLUP-AD model are smaller than GBLUP-A, ABLUP-AD, and ABLUP-A models, which indicates that GBLUP-AD could produce more accurate breeding values even though predictive accuracy or Spearman breeding values ranking correlation are similar between GBLUP-AD and GBLUP-A.

### M×E for genomic selection in multi-environment trials (MET)

Gamal EL-Dien et al. (2018) showed that interior spruce (*Picea glauca* x *engelmannii*) had substantially significant additive M×E and non-significant small dominance M×E terms for both height and wood density. In our study, significant, but small M×E effects for all traits were only found in additive genetic effects, not for dominance. Gamal EL-Dien et al. (2018) did not use more comprehensive models to dissect M×E, but used compound symmetry variance-covariance structure (CS). To more accurately dissect M×E in multi-environment trials (MET), here we used six variance-covariance matrices (Table 1) to model additive and dominance effects in GS models for four traits. A similar approach was described by Burgueño et al. (2012) and Oakey et al. (2016) and Ventorim Eerrao et al. (2017). Finally, we find that all four traits have significant additive M×E term using CS for additive effects. For tree height, we also observe a better goodness-of-fit using compound symmetry with heterogeneous variance (CS+DIAG) (Cullis et al.1998), unstructured (US), and factor analytic (FAMK) (Smith et al. 2001) in Table S1. Here we should note that site 1 has a non-significant additive M×E term for tree height that resulted in a negligible increase with the M×E term included in GBLUP-AD model. Generally, MET analysis shows slightly higher PA than ST analysis, except for tree height with the same value. It may result from the non-significant additive covariance 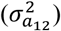 between two sites. In this study, only tree height and velocity had slight increases for PA, which also supports the previous study of Ly et al. (2013) that including the G×E term could improve the PA.

### Significant dominance effects improve predictive ability

Recent studies have shown that maximum PA can be reached when the model was based on additive and non-additive effects (Aliloo et al. 2016; Da et al. 2014; Muñoz et al. 2014; Tan et al. 2018). Ly et al. (2013) considered that only the additive component may produce a systematic underestimation of PA because only additive effects are predicted. Here, the GBLUP-AD model for tree height shows homogenous dominance variances in both sites (Table 3, identity matrix for dominance effect). However, the ABLUP-AD model shows that a significant dominance variance in site 2 (23.1%) and non-significant dominance variance in site 1 (8.5%), indicating GBLUP-AD model has higher efficiency in separating the additive and dominance genetic variances because it could account for the Mendelian sampling within families for dominance.

It was found that including dominance could improve PA when a considerable dominance variance in animal (Aliloo et al. 2016; Esfandyari et al. 2016) and plants studies were observed (Tan et al. 2018; Wolfe et al. 2016). In this study, the improvement of tree height and velocity PAs also agree with the previous observations. However, including significant dominance in this study may not improve the additive accuracy (Table 3, the diagonal values) and breeding values ranking correlation (Table 4, the diagonal values).

In several practical breeding programs, a dominance effect has been utilized, such as Sitka spruce (*Picea sitchensis*) (Thompson, 2013), loblolly pine (McKeand et al. 2003), and Eucalypts (McKeand et al. 2003; Rezende et al. 2014). For instance, since 2000, the annual production of full-sib seedlings in loblolly pine has increased to 63.2 million seedlings in 2013, with a total of over 325 million full-sib family seedlings planted over last 14 year (Steve Mckeand 2014, personal communication). In Norway spruce, dominance estimate was not widely included in the breeding program, but we are commonly using full-sib family materials. Thus, it is important to estimate dominance effects in Norway spruce breeding program when more and more individuals will be genotyped for selection and propagation.

### Epistasis effect

The full model (GBLUP-ADE), which was extended to include three first-order interactions, shows almost the same results as GBLUP-AD for all four traits based on AIC (Table 2). This indicates the absence of three kinds of epistatic interactions even though additive × additive and dominance × dominance epistatic effects explained variations of 4.2% (4.0%) and 0.7% (0.2%) for velocity in site 1 (site2), respectively. However, in several other forest tree species, such as white spruce (Gamal El-Dien et al. 2016), interior spruce (*P. glauca x engelmannii*) (Gamal El-Dien et al. 2018), and loblolly pine (de Almeida Filho et al. 2016) and Eucalypt (Bouvet et al. 2016; Resende et al. 2017; Tan et al. 2018) significant epistatic effects have been reported for height or wood density. For instance, Gamal El-Dien et al. (2016) showed a significant additive × additive component and non-significant dominance for tree height, while Gamal El-Dien et al. (2018) showed a considerable dominance component (19.46% of total phenotypic variation) and none epistatic effect. For wood density in spruce, Gamal El-Dien et al. (2016, 2018) showed that a significant additive × additive interaction that was absorbed from additive and residuals variances. Tan et al. (2018) showed none of the epistasis for wood density. Above results agree with the suggestion by Tan et al. (2018) that the contributions of non-additive effects, especially epistasis effects, are traits, populations, and species-specific, even site-specific as in this study. However, including non-additive effects could improve the estimates of genetic parameters when they are significant.

### Expected response to genomic selection

The main advantages of GS are to shorten the length of the breeding cycle and, reduce phenotypic evaluation costs in plant and animal breeding (de los Campos et al. 2013; Grattapaglia and Resende, 2011). In Northern Sweden, the breeding cycle length of Norway spruce in GS could be ideally shortened from 30 years to 15 years (Chen et al. 2018) if we could complete flowering induction and conducting controlled pollination within 15 years. In our previous paper (Chen et al. 2018), we calculated the RGS per year for GS based on GBLUP-A using the same data set. Here we compared RPS with RGS per year for GS based on GBLUP-AD model and calculated the response to selection per year for PS and GS. We used EGVs from ABLUP-AD model as a benchmark for all traits. RGS per year is considerably higher than RPS per year for all traits (Figure 3). RGS per year for wood quality traits has higher gain than those for tree height when we select the top 50 individuals based on M, A or AD effect, in contrast to the result reported by Resende et al. (2017) at *Eucalyptus*. Thus, GS based on GEGV is ideal for solid-wood quality improvement in Norway spruce.

## Conclusions

This is the first paper to study M×E using different covariance structure for the additive and non-additive effects and dominance in GS for forestry trees species. We found that M×E and dominance effects could improve PA when they are considerably large. In GBLUP-AD model, M×E contributed 4.7% and 11.1% of tree height phenotypic variations for the site1 and site 2, respectively. Dominance contributed 18.1% and 9.8% of tree height phenotypic variations for the site1 and site 2, respectively. The higher PA of GBLUP-AD model for tree height compared to ABLUP-A and GBLUP-A models suggests that dominance should be included in GS models for the forestry genetic evaluation in order to improve the predictive accuracy or estimates of genetic parameters. Advanced M×E model could improve PA and should be kept in the model fitting. GBLUP-AD could be a more useful model in breeding and propagation when tree breeders want to use the specific combining ability (SCA) using full-sib family seedlings.

## Data archiving

The Data will be archived after the manuscript is accepted.

## Acknowledgements

Financial support was received from Formas (grant number 230-2014-427) and the Swedish Foundation for Strategic Research (SSF, grant number RBP14-0040). The computations were performed on resources by the Swedish National Infrastructure for Computing (SNIC) at UPPMAX and HPC2N. We thank Dr. Junjie Zhang, Tianyi Liu, and Ms Xinyu Chen, Linghua Zhou, Ruiqi Pian for help of the DNA extraction and field assistance and Anders Fries for field work.

